# Recent Endemic Coronavirus Infection Associates With Higher SARS-CoV-2 Cross-Reactive Fc Receptor Binding Antibodies

**DOI:** 10.1101/2024.10.23.619886

**Authors:** David J. Bean, Yan Mei Liang, Manish Sagar

**Affiliations:** Department of Virology, Immunology and Microbiology, Boston University Chobanian & Avedisian School of Medicine; Boston, MA; Department of Medicine, Boston University Chobanian & Avedisian School of Medicine; Boston, MA

## Abstract

Recent documented infection with an endemic coronavirus (eCoV) associates with less severe coronavirus disease 2019 (COVID-19), yet the immune mechanism behind this protection has not been fully explored. We measured both antibody and T cell responses against severe acute respiratory syndrome coronavirus 2 (SARS-CoV-2) in SARS-CoV-2 naïve individuals classified into two groups, either with or without presumed recent eCoV infections. There was no difference in neutralizing antibodies and T cell responses against SARS-CoV-2 antigens between the two groups. SARS-CoV-2 naïve individuals with recent presumed eCoV infection, however, had higher levels of Fc receptor (FcR) binding antibodies against eCoV spikes (S) and SARS-CoV-2 S2. There was also a significant correlation between eCoV and SARS-CoV-2 FcR binding antibodies. Recent eCoV infection boosts cross-reactive antibodies that can mediate Fc effector functions, and this may play a role in the observed heterotypic immune protection against severe COVID-19.

## INTRODUCTION

Prior to the emergence of severe acute respiratory syndrome coronavirus 2 (SARS-CoV-2) and coronavirus disease 2019 (COVID-19), studies suggested that humans are infected with an endemic coronavirus (eCoV), such as human coronavirus (HCoV)-229E, HCoV-HKU1, HCoV-NL63, and HCoV-OC43, about every 18 months to 2 years (1, 2). It has been hypothesized that cross-reactive immune responses generated from one coronavirus (CoV) may protect against infection or ameliorate disease severity from a heterologous CoV (3, 4). In a large retrospective cohort study, we observed that individuals with a documented recent eCoV infection had less severe COVID-19 although there was no difference in SARS-CoV-2 incidence (5). The immune mechanism behind this observed protection has not been definitively elucidated. Some, but not all, studies suggest that the pre-existing humoral and cellular immunity may decrease SARS-CoV-2 incidence and COVID-19 severity (6–8). Few studies, however, have directly looked at how the cross-reactive immune response changes after a recent eCoV infection.

Recent eCoV infections are difficult to diagnose because disease is mild, and the etiology for a “common cold” is usually not investigated if one presents for medical care (9). Many studies define recent eCoV infections either by detecting viral RNA through polymerase chain reaction (PCR)-based methods or changes in eCoV directed antibody levels (2, 10). Comprehensive respiratory panel (CRP)-PCR (BioFire) tests detect active eCoV infections (11), but they are not widely used and most individuals do not present for medical care when they have the “common cold” due to an eCoV (10). Detecting changes in antibody levels require longitudinal sampling, and these types of studies are hindered by the number of participants and longevity of the study. Inability to identify individuals with recent eCoV infections with confidence makes it difficult to assess the effect on SARS-CoV-2 immunity.

In this study, we used a combination of PCR documented eCoV infections with eCoV nucleocapsid antibody titer data to classify individuals with presumed recent eCoV infections. The cross-reactive immune responses to various SARS-CoV-2 antigens were compared between those individuals with or without a presumed recent eCoV infection. We found that eCoV spike specific antibodies are boosted after a recent eCoV infection and are likely mediating the cross-reactivity against SARS-CoV-2 spike. These cross-reactive antibodies were not associated with higher SARS-CoV-2 neutralization, but they were capable of binding to the Fc receptor (FcR), FcγRIIIa, and potentially mediating enhanced Fc effector functions. Our results suggest a recent infection with an eCoV boosts the Fc effector function potential of CoV-specific antibodies and offers an additional possible mechanism for the protection against severe COVID-19.

## RESULTS

### Plasma IgG levels indicate recent CoV infections

We examined immune responses in blood samples from individuals that either had a confirmed previous SARS-CoV-2 infection (n=20), prior COVID-19 vaccination (n=29), or no known history of SARS-CoV-2 infection or COVID-19 vaccination (n=72) (Table S1). Of those with no documented SARS-CoV-2 infection or COVID-19 vaccination, 18 were sampled prior to the start of the COVID-19 pandemic in the United States. Thus, these individuals could not have had undocumented or asymptomatic SARS-CoV-2 infection. The remaining people (n=54), however, possibly could have had prior undiagnosed or asymptomatic SARS-CoV-2 infection because samples were collected after March 2020. We used previous methodology described by our group and others to identify individuals that may have had undocumented asymptomatic SARS-CoV-2 infection (4, 12). Briefly, we measured plasma IgG levels against SARS-CoV-2 receptor binding domain (RBD) and SARS-CoV-2 nucleocapsid by enzyme-linked immunosorbent assay (ELISA). These ELISA results on samples collected prior to the pandemic, those with documented SARS-CoV-2 infection, and those with known COVID-19 vaccination were used to set cutoffs for the SARS-CoV-2 RBD and nucleocapsid IgG levels that would differentiate individuals with possible asymptomatic undocumented SARS-CoV-2 infections (Figure 1A). As expected, most of the individuals with either a previous SARS-CoV-2 infection (18/20, 90%) or COVID-19 vaccination (28/29, 97%) had elevated anti-SARS-CoV-2 RBD IgG titers. Furthermore, all the pre-pandemic samples (18/18, 100%) were below the set cutoff for SARS-CoV-2 antigen exposure. In summary, the overall classification accuracy based on our assigned cutoffs was 87% (95% confidence interval (CI) 76–94%), which is within the range of previous methodologies (73 to 91%) (Figure 1B) (4). Seven individuals classified in the no known SARS-CoV-2 infection or COVID-19 vaccination group had IgG levels against SARS-CoV-2 RBD above the established cutoff, indicative of a potential previous SARS-CoV-2 infection. Furthermore, anti-SARS-CoV-2 nucleocapsid IgG levels above the established cutoff indicated that 6 individuals with a prior COVID-19 vaccination may have had a previous SARS-CoV-2 infection. These 13 individuals were excluded from the subsequent groups classified according to possible recent eCoV infection.

**Figure 1.**
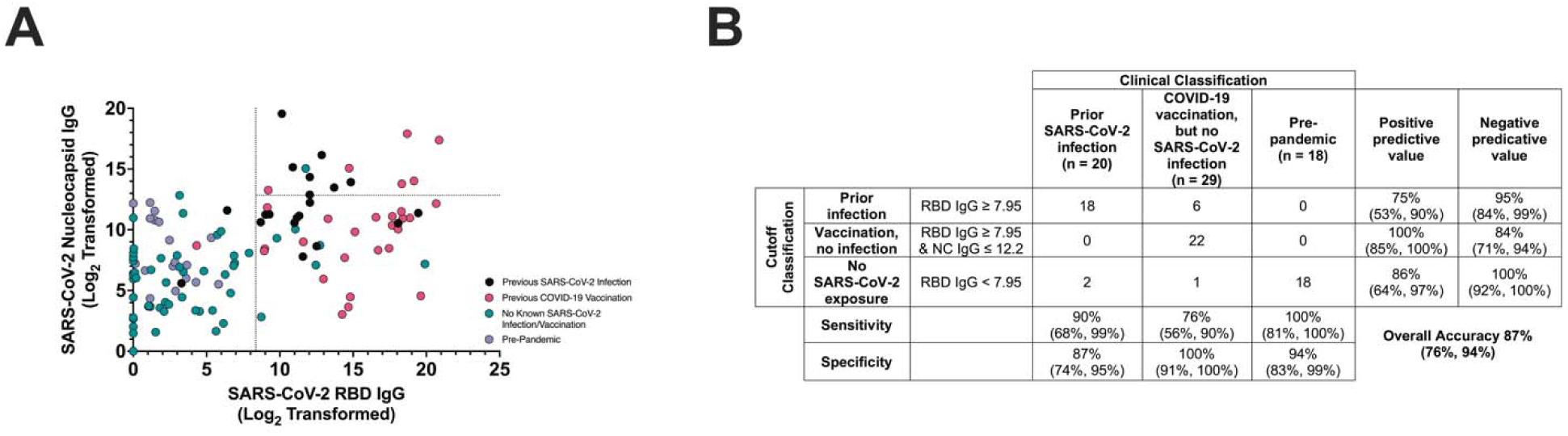
Classification of presumed SARS-CoV-2 infections based on antibody levels. (A) IgG antibody levels against SARS-CoV-2 RBD and nucleocapsid protein were measured in individuals with a previous SARS-CoV-2 infection (black), previous COVID-19 vaccination with no documented history of SARS-CoV-2 infections (pink), no known SARS-CoV-2 infection or COVID-19 vaccination (green), or pre-pandemic samples collected before March 2020 (purple). The dotted lines specify cutoffs that were established to classify individuals with presumed undiagnosed SARS-CoV-2 infection. (B) The accuracy and other test characteristics of the classification based on the established cutoffs for classifying individuals into the specified groups. The values in parentheses display the 95% CI.

Similarly, plasma IgG antibody levels against the nucleocapsid antigen have been previously used to identify recent eCoV infections (13). We measured plasma IgG levels against all four human eCoVs nucleocapsid protein to identify presumed recent eCoV infections within our COVID-19 vaccinated and no known SARS-CoV-2 exposure groups. Nucleocapsid antibody titers against the two alpha (HCoV-229E and HCoV-NL63, Pearson ρ=0.920, p<0.0001, Fig. S1A) and the two beta (HCoV-OC43 and HCoV-HKU1, Pearson ρ=0.478, p<0.0001, Fig. S1B) eCoVs showed significant correlations. On the other hand, there was no correlation between alpha and beta nucleocapsid (Figure S1C). Thus, we generated a composite alpha and beta eCoV anti-nucleocapsid IgG metric. Longitudinal sampling in prior studies suggests a reinfection with an eCoV occurs on average every 1 to 3 years (1, 2). Furthermore, duration between peak antibody responses after a presumed eCoV infection suggest that a subsequent infection with a heterologous eCoV occurs around every 1.5 years (2).

Based off this timeframe and the Boston Medical Center (BMC) electronic medical record (EMR), we identified 22 individuals within the no known SARS-CoV-2 infection that had documented eCoV RNA (11 HCoV-OC43, 6 HCoV-NL63, 4 HCoV-229E, 1 HCoV-HKU1) on a prior CRP-PCR test less than 550 days prior to sample collection. Thus, these individuals were definitively classified as having a recent eCoV infection. In the remaining with no documented or presumed SARS-CoV-2 infection (n=72), the top 25% of individuals (n=18) with the highest nucleocapsid antibody levels (either anti-alpha or beta eCoV) were also classified as having a recent eCoV infection. The rest (n=54) were classified as not having a recent eCoV infection Other than a clinically small but statistically significant age difference, the groups classified as either with or without a recent eCoV infection had no meaningful demographic differences (Table 1).

### Recent eCoV infection associates with increased SARS-CoV-2 S2 specific FcγRIIIa binding antibody responses

We examined cross-reactive humoral immunity after a recent eCoV infection because antibody responses are the best correlate of protection against SARS-CoV-2 infection and severe COVID-19 (14, 15). Individuals with a previous COVID-19 vaccination were excluded from the groups classified as with (n=13) or without (n=10) a presumed recent eCoV infection when comparing heterotypic SARS-CoV-2 spike immune responses (Table 1). Instead, they were included in a composite group along with those with a documented previous SARS-CoV-2 infection, classified as previous SARS-CoV-2 spike exposure. There was no significant difference in anti-SARS-CoV-2 RBD IgG (Figure 2A) and anti-SARS-CoV-2 S2 (Figure 2B) IgG antibodies in those with and without a recent eCoV infection. As expected, anti-SARS-CoV-2 spike antibodies were significantly higher in the individuals with prior SARS-CoV-2 spike exposure (Figure 2A and B). Neutralization responses against vesicular stomatitis virus (VSV) pseudotyped with SARS-CoV-2 spike (Wuhan variant) were higher in those with previous SARS-CoV-2 spike exposure, but there was no difference in neutralization between individuals with or without a presumed recent eCoV infection (Figure 2C).

**Figure 2.**
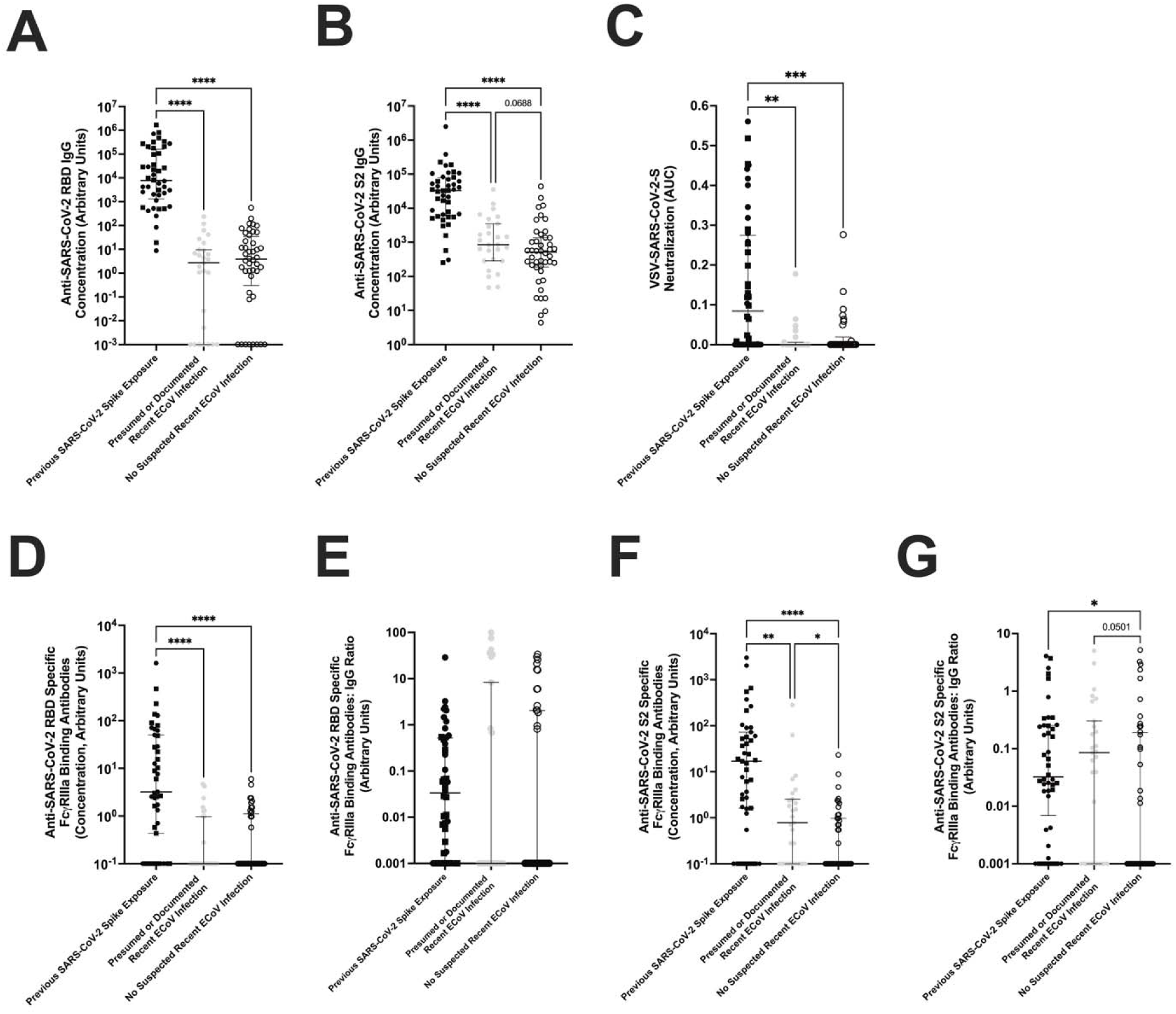
Humoral immune responses to SARS-CoV-2 spike antigens. Plasma antibody responses in those with documented SARS-CoV-2 spike exposure (previous SARS-CoV-2 infection (black circle) or prior COVID-19 vaccination (black square)), presumed or documented recent eCoV infection (gray circle), or without a presumed recent eCoV infection (white circle). (A and B) IgG antibody levels against SARS-CoV-2 RBD (A) or SARS-CoV-2 spike S2 subunit (B). (C) Neutralization responses against a VSV-ΔG pseudovirus expressing the SARS-CoV-2-Wuhan spike protein. (D-G) Titer of SARS-CoV-2 RBD (D) or SARS-CoV-2 S2 (F) specific antibodies binding to the Fc receptor, FcγRIIIa, or the ratio of those FcR binding antibodies to the SARS-CoV-2 RBD (E) or SARS-CoV-2 S2 (G) IgG levels in panel (A and B). The dark horizontal lines in each scatter dot plot denote the median and interquartile range. Statistical analyses were performed using either Kruskal-Wallis test with Dunn multiple comparison test or Mann-Whitney U test. P-values less than 0.1 are displayed. *, **, ***, and **** represent p-values <0.05, <0.01, <0.001, and <0.0001, respectively.

Another potential functional role of these anti-SARS-CoV-2 antibodies is through FcR binding, which contributes to the clearance of virally infected cells through mechanisms such as antibody-dependent cellular cytotoxicity (ADCC) or antibody-dependent cellular phagocytosis (ADCP) (16). We used a previously validated ELISA method (17) to measure SARS-CoV-2 specific FcγRIIIa (CD16a) binding antibodies, which is a predictor for natural killer (NK) cell mediated antibody effector function. As expected, those individuals with prior SARS-CoV-2 spike exposure had higher levels of anti-SARS-CoV-2 RBD and S2 antibodies binding to FcγRIIIa (Figure 2D-G). Individuals with or without a presumed recent eCoV infection had similar anti-SARS-CoV-2 RBD FcγRIIIa binding antibodies and FcγRIIIa binding antibodies to IgG ratio (Figure 2D and E). However, anti-SARS-CoV-2 S2 FcγRIIIa binding antibodies were 7.8-fold higher (p=0.0234) and the FcγRIIIa binding antibodies to IgG ratio was 85-fold higher (p=0.0501) in the individuals classified as having a recent eCoV infection (Figure 2F and G).

In the FcγRIIIa binding assay, individuals could be separated into responders and non-responders based on FcγRIIIa binding activity above or below a designated cutoff (Figure 3A). We used a multiple logistic regression model to identify characteristics associated with SARS-CoV-2 S2-specific FcγRIIIa antibody binding. As expected, previous SARS-CoV-2 spike exposure was associated with 4.055-fold higher odds (95% CI: 1.581 to 11.55, p=0.0053) of having a SARS-CoV-2 S2 FcγRIIIa binding antibody response. There were 1.379-fold higher odds (95% CI: 1.060 to 1.859, p=0.0249) of having a SARS-CoV-2 S2 FcγRIIIa binding antibody response for every 2-fold increase in anti-alpha eCoV nucleocapsid antibody levels (Figure 3B). Levels of anti-beta eCoV nucleocapsid antibodies, age, gender, and other demographic characteristics were not associated with detectable SARS-CoV-2 S2 FcγRIIIa binding antibody.

**Figure 3.**
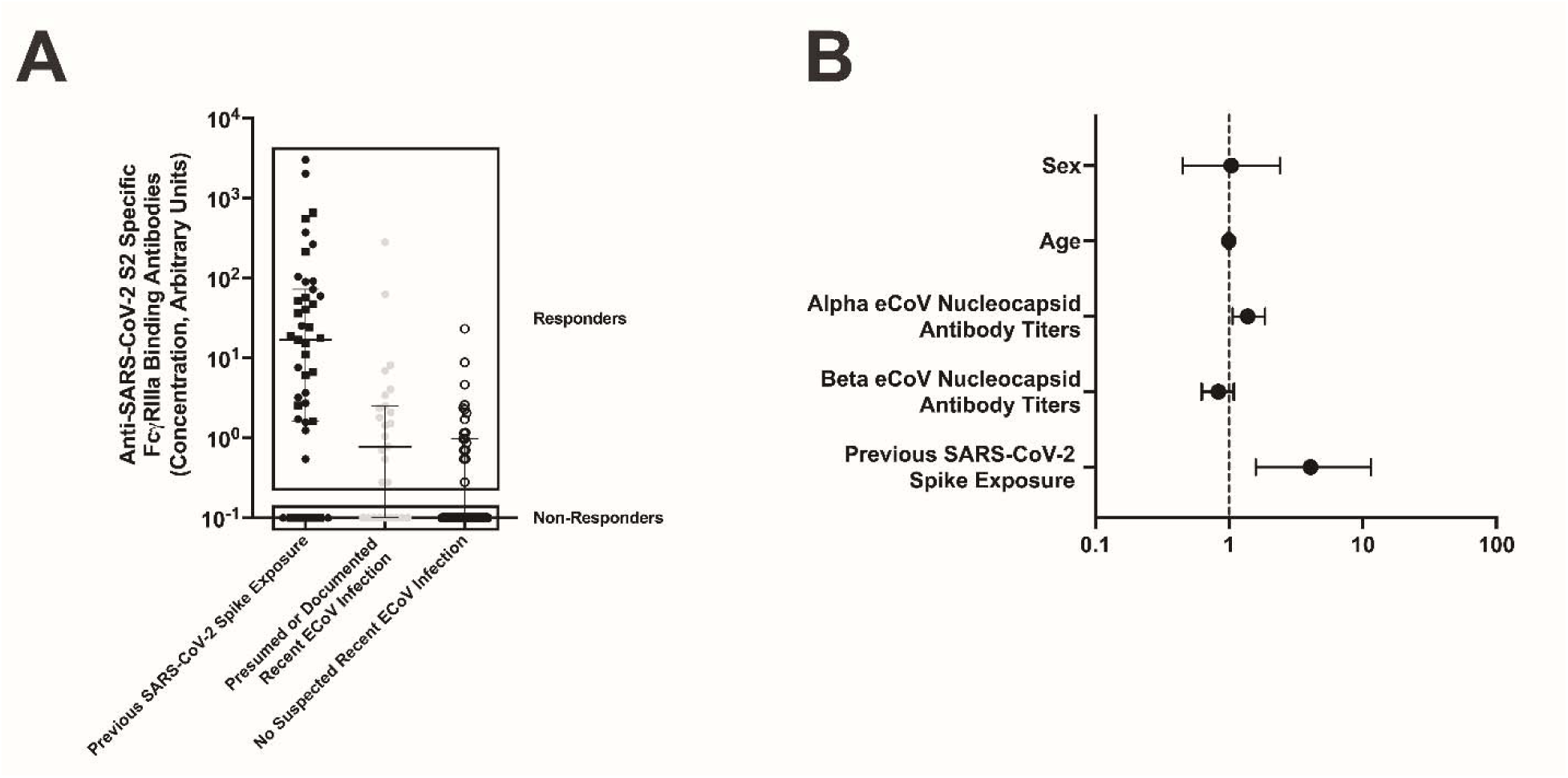
Higher alpha eCoV nucleocapsid antibody responses predicts positive responses in the SARS-CoV-2 S2 specific antibody FcR binding assay. (A) Individuals were separated into two groups based on positive (responders) or absent (non-responders) FcγRIIIa SARS-CoV-2 S2 specific antibody binding. This responder and non-responder classification was used in a multivariate logistic regression model. This data is the same as Fig. 2F. (B) The odds ratio of variables used in the multivariate logistic regression model to predict positive SARS-CoV-2 S2 antibody FcR binding responses. The error bars represent the 95% CI.

### SARS-CoV-2 S2 Fc**γ**RIIIa binding antibody responses are correlated with eCoV spike specific Fc**γ**RIIIa binding antibody responses

Next, we assessed if eCoV spike protein directed FcγRIIIa binding antibodies possibly account for the anti-SARS-CoV-2 S2 FcγRIIIa antibodies (Figure 2 and 3). We used the antigen specific IgG and FcγRIIIa binding antibody ELISA to measure antibody levels against HCoV-229E spike and HCoV-OC43 spike proteins as representative antibody responses to alpha and beta eCoV infections respectively (Figure 4). As expected, IgG levels were higher against HCoV-229E spike (p=0.0241, Figure 4A) and HCoV-OC43 spike (p=0.2086, Figure 4B) in the individuals with a presumed recent eCoV infection, although only the comparison with HCoV-229E spike antibody levels reached statistical significance. Furthermore, HCoV-229E (p=0.0688, Figure 4C) and HCoV-OC43 (p=0.0282, Figure 4D) spike specific FcγRIIIa binding antibody levels were also higher in the individuals with a presumed recent eCoV infection, and these comparisons showed a statistical trend and significance. There were no statistical differences in the HCoV-229E (p=0.4689, Figure 4F) and HCoV-OC43 spike specific (p=0.2044, Figure 4E) FcγRIIIa binding antibodies to IgG ratio between those with or without a presumed recent eCoV infection.

**Figure 4.**
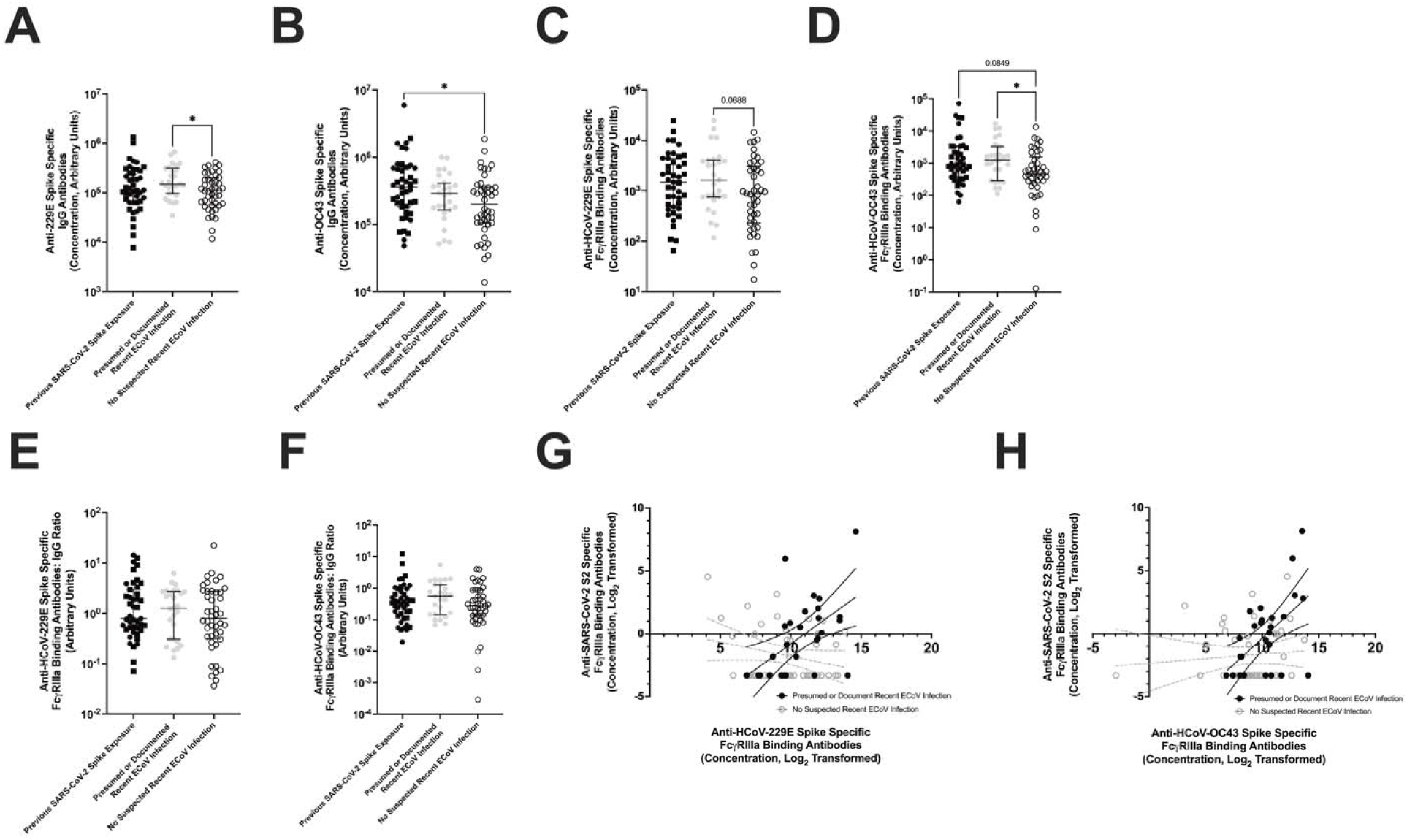
ECoV specific antibody FcR binding responses correlate with SARS-CoV-2 S2 antibody responses. Plasma antibody responses were measured in individuals with either a previous SARS-CoV-2 spike exposure (previous SARS-CoV-2 infection (black circle) and previous COVID-19 vaccination (black square)), presumed or documented recent eCoV infection (gray circle), or without a presumed recent eCoV infection (white circle). (A and B) IgG antibody responses against HCoV-229E (A) or HCoV-OC43 (B) spikes. (C and D) Level of HCoV-229E (C) or HCoV-OC43 (D) spike-specific antibodies binding to the Fc receptor, FcγRIIIa. (E and F) Ratio of Fc receptor antibody binding (C and D) to antigen specific IgG (A and B) in HCoV-229E (E) or HCoV-OC43 (F) spike-specific antibody responses. (G and H) Correlations between HCoV-229E (E) or HCoV-OC43 (F) spike-specific antibody FcR binding to SARS-CoV-2 S2 specific antibody FcR binding responses. Black dots and lines represent individuals with presumed or documented recent eCoV infection, while gray represents those without a presumed recent eCoV infection. The lines show a simple linear regression with the 95% CI. The dark horizontal lines in each scatter dot plot denote the median and interquartile range. Statistical analyses were performed using either Kruskal-Wallis test with Dunn multiple comparison test or Mann-Whitney U test. P-values less than 0.1 are displayed. * represents p-values <0.05.

We compared the SARS-CoV-2 S2 and eCoV spike FcγRIIIa binding antibody responses to better understand the extent of cross-reactivity of the antibodies. Moderate positive correlations were observed between the SARS-CoV-2 S2 and HCoV-229E (Spearman r=0.5153 and p=0.010, Figure 4G) and HCoV-OC43 (Spearman r=0.4793 and p=0.010 Figure 4H) spike FcγRIIIa binding antibody responses in those individuals with presumed or documented recent eCoV infections. There were no significant correlations in SARS-CoV-2 and eCoV FcR binding antibody responses in individuals without a presumed recent eCoV infection. These observations suggest eCoV antibodies spike capable of binding to FcγRIIIa are elevated after a recent eCoV infection, and these antibodies correlate with increased cross-reactivity against SARS-CoV-2 spike S2 region.

### Recent eCoV infection does not improve T cell responses against SARS-CoV-2 antigens

We have previously shown that T cell responses generated by SARS-CoV-2 infection associate with lower incidence of eCoV infections, which demonstrated the impact of heterotypic T cell responses among the different CoVs (4). We tested T cell responses against SARS-CoV-2 spike, nucleocapsid, and nsp12/nsp13 antigens in the subset of individuals with available peripheral blood mononuclear cells (PBMC) and either a confirmed previous SARS-CoV-2 infection (n=20), presumed or documented recent eCoV infection (n=32), or no presumed recent eCoV infection (n=42). The subset of individuals with available PBMCs were representative of the full cohort (Table S3). As before, individuals with a previous COVID-19 vaccination were included in the prior SARS-CoV-2 spike exposure group when testing responses against SARS-CoV-2 spike. As expected, individuals with a previous SARS-CoV-2 antigen experience had significantly higher activated (CD134^+^ CD137^+^) CD4^+^ and (CD69^+^ CD137^+^) CD8^+^ T cell responses to SARS-CoV-2 spike (Fig. 5A and B) and nucleocapsid (Fig. 5C and 5D).

**Figure 5.**
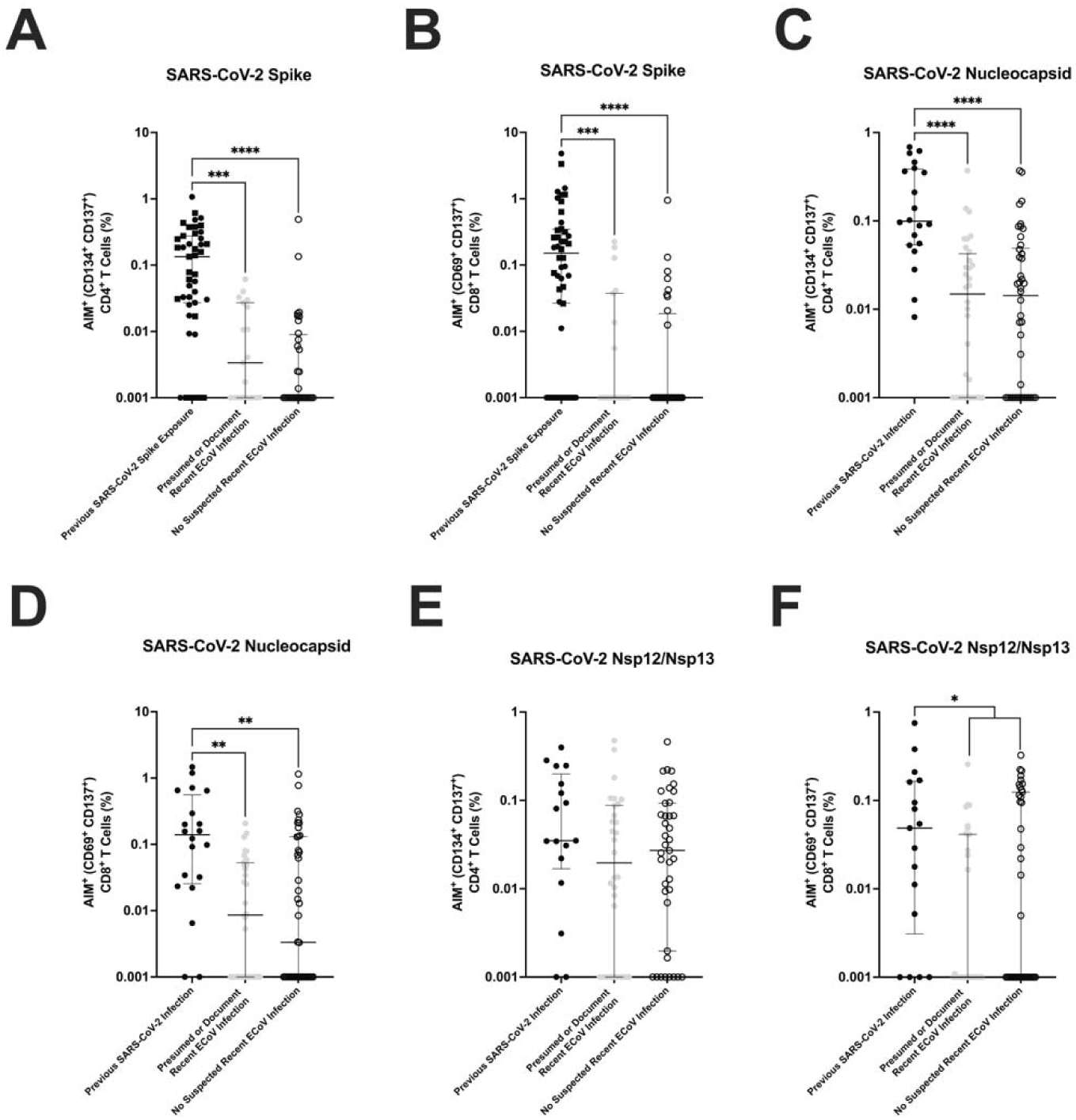
T cell responses against SARS-CoV-2 antigens are similar between individuals with or without presumed recent eCoV infections. T cell responses were measured in individuals with either a previous SARS-CoV-2 spike exposure (previous SARS-CoV-2 infection (black circle) and previous COVID-19 vaccination (black square)), presumed or documented recent eCoV infection (gray circle), or without a presumed recent eCoV infection (white circle). Cells were stimulated with SARS-CoV-2 peptide pools and percent of activated (CD134^+^ CD137^+^) CD4^+^ T cells and (CD69^+^ CD137^+^) CD8^+^ T cells were measured. (A-B) CD4^+^ (A) and CD8^+^ (B) T cell activation was measured after stimulation with SARS-CoV-2 spike peptides. (C-D) CD4^+^ (C) and CD8^+^ (D) T cell activation was measured after stimulation with SARS-CoV-2 nucleocapsid peptides. (E-F) CD4^+^ (E) and CD8^+^ (F) T cell activation was measured after stimulation with SARS-CoV-2 nsp12/nsp13 peptides. Data were background subtracted against the negative control (DMSO only). The dark horizontal lines in each scatter dot plot denote the median and interquartile range. Note, the y axis varies among the different panels. Statistical analyses were performed using either Kruskal-Wallis test with Dunn multiple comparison test or Mann-Whitney U test. P-values less than 0.1 are displayed. *, **, ***, and **** represent p-values <0.05, <0.01, <0.001, and <0.0001, respectively.

Nsp12/nsp13 CD4^+^ (p=0.1120, Fig. 5E) and CD8^+^ T cell responses (p=0.0327, Fig. 5F) were also higher in those with prior SARS-CoV-2 infection as compared to those with no previous SARS-CoV-2 antigen experience, although there were no consistent differences when compared with the two groups classified based on recent eCoV infection individually. Importantly, there were no significant differences in activated CD4^+^ or CD8^+^ T cell responses to SARS-CoV-2 spike, nucleocapsid, or nsp12/nsp13 peptides between those individuals with or without a presumed recent eCoV infection (Figure 5A-F). This suggests that cross-reactive T cell levels in the peripheral blood are not significantly boosted after a recent eCoV infection.

## DISCUSSION

Multiple different highly pathogenic CoV have emerged over the past 25 years (18–20), and thus developing a pan-CoV vaccine remains a major priority. The efficacy of this pan-CoV vaccine against future emerging CoVs may be judged by how well it performs against the known currently circulating HCoVs. Such broad protection against the diverse CoVs requires knowledge of both the conserved immunodominant regions and the components of the immune system mediating this protection. Understanding the role of heterotypic immunity in the protection against SARS-CoV-2 and severe COVID-19 is a key step towards developing a pan-CoV vaccine. Most studies suggest that heterotypic immunity from a recent eCoV infection protects against severe COVID-19, while some studies indicate there is no cross-reactive protection (5, 21–23). These varying conclusions may differ based on the cohorts studied and criteria used to define protection against SARS-CoV-2 or severe COVID-19. Besides resolving this controversy, it is important to understand how a recent eCoV infection impacts immune responses directed against SARS-CoV-2 because it will highlight immune mechanisms that mediate cross-reactive protection. Here in this study, we identify a potential role of antibody Fc effector functions in the eCoV-derived heterotypic immune response against SARS-CoV-2.

Studies have measured heterotypic immunity against SARS-CoV-2 by correlating eCoV and SARS-CoV-2 immune responses (24, 25). In general, investigations have not identified individuals with a recent eCoV infection and assessed subsequent SARS-CoV-2 immune responses. We used both clinically documented cases and antibody levels against eCoV nucleocapsids to classify presumed recent eCoV infections. These are the most reliable and accepted ways to classify recent eCoV infection. This classification, however, is complicated by the prevalence and frequency of eCoV infections. By the age of 5, most children have had at least 1 eCoV infection with reinfections occurring frequently (10, 26). The protective immunity towards these eCoVs lasts for a short duration, and reinfections can even happen within the same year despite detectable antibody levels (2). Thus, it is difficult to accurately define a recent eCoV infection without complete PCR-based results or longitudinal sampling of the entire cohort. Our results and comparisons between groups largely depend on this definition of a recent eCoV infection. SARS-CoV-2 directed FcR antibodies still trended higher if the prior eCoV infection group was defined by documented infection on CRP-PCR test within the past 550 days and up to 35% of those with the highest alpha and beta nucleocapsid IgG levels (Table S3). This sensitivity analysis provides further confidence for our conclusions. Regardless, additional studies with well-defined eCoV infections will further validate our findings.

In our study, SARS-CoV-2 S2, but not RBD, directed antibodies were elevated after a recent eCoV infection (Figure 2). These results align with previous studies because many individuals have pre-existing anti-SARS-CoV-2 antibodies, with a preference towards more conserved regions, like S2, compared to more variable regions, like RBD (27). In general, the higher degree of similarity between different CoVs predicts higher levels of antibody-mediated protection against heterologous infections (3). Sterilizing protection against a heterologous CoV requires neutralizing antibodies against conserved spike regions. In accordance with our study and previous work, eCoV directed antibodies poorly cross-react with SARS-CoV-2 RBD and thus have limited neutralizing ability (22). Antibodies targeted against SARS-CoV-2 S2 can neutralize the virus, but they are rare and less potent than anti-SARS-CoV-2 RBD neutralizing antibodies (28). This agrees with clinical observations that a recent eCoV infection does not reduce the number of SARS-CoV-2 infections, but instead associates with protection against COVID-19 severity (5, 21). The protection against severe COVID-19, but not SARS-CoV-2 infection, indicates components of the immune system other than neutralizing antibodies may be mediating this protection.

The Fc effector function of cross-reactive SARS-CoV-2 directed antibodies could potentially contribute to the protection against severe COVID-19 (29). SARS-CoV-2 S2 FcγRIIIa binding antibodies were higher in those classified as having a recent eCoV infection, increased levels were associated with higher alpha eCoV nucleocapsid levels, and there was a direct association with eCoV spike IgG levels. In aggregate, this implies that recent eCoV infection enhances cross-reactive SARS-CoV-2 S2 FcγRIIIa binding antibodies. We used an ELISA which measures antigen specific antibodies capable of binding to FcγRIIIa, which primarily depends of afucosylation of the antibody Fc portion (17). We focused our study on FcγRIIIa, which is expressed on natural killer (NK) cells and macrophages, and its interaction is crucial for ADCC and ADCP activity (30). Other Fc receptors, like FcγRI, have a higher affinity toward IgG antibodies, but they are not expressed on NK cells and do not play as critical of a role in Fc mediated effector functions (31). We found that both SARS-CoV-2 S2 and eCoV spike antibodies capable of binding FcγRIIIa were increased in those individuals with a recent eCoV infection.

Both higher afucosylated antibody levels and greater ADCC activity are correlated with COVID-19 severity (32, 33). Yet, stronger ADCC activity is also associated with protection against fatal COVID-19 cases (16). The Fc portion of antibodies can also enhance viral infections and contribute to disease pathogenesis. Antibody dependent enhancement (ADE) may increase SARS-CoV-2 infection in monocytes, but its role in COVID-19 severity is not clear (34). We only measured levels and not the function of FcγRIIIa binding antibodies, and thus we can only speculate about increases in ADCC, ADCP, and ADE. Furthermore, we restricted our analysis to peripheral blood immune responses, and cross-reactive tissue-based SARS-CoV-2 immunity may be different after recent eCoV infection Elevated levels of T cells early in a SARS-CoV-2 infection are associated with protection against severe COVID-19 (35, 36). These initial protective T cells against SARS-CoV-2 are speculated to be cross-reactive memory T cells from a previous eCoV infection. Although other studies have noted a boost in cross-reactive T cells after a SARS-CoV-2 infection, few studies have looked at cross-reactive T cell levels after an eCoV infection (4, 37). Cross-reactive T cells against SARS-CoV-2 antigens were detected in nearly all SARS-CoV-2 naïve individuals, but they were at similar levels among those with or without a presumed recent eCoV infection. Our data suggests that an eCoV infection does not preferentially expand T cell responses against SARS-CoV-2 spike, nucleocapsid, or nsp12/nsp13. Alternatively, eCoV-specific T cells have a lower avidity toward their corresponding SARS-CoV-2 peptides, so the cross-reactive T cells may not respond sufficiently in our assay (38). Furthermore, we only stimulated cells with peptides from SARS-CoV-2 spike, nucleocapsid, nsp12, and nsp13. We chose these proteins because they are either immunodominant or associated with immune protection against CoV related diseases, but other antigens may contribute to the heterotypic immune response (4, 6, 39). Humans have a limited T cell repertoire against SARS-CoV-2 epitopes, and thus it is possible that the appropriate peptides were not included in our peptide pools (39). Cross-reactive T cells may still contribute to reducing COVID-19 severity, but in our cohort, they were not associated with a recent eCoV infection.

Our observations suggest that antibody Fc effector function may play a critical role in the reduction of COVID-19 severity. Antibody Fc effector functions are important in the battle against other viruses and disease, including HIV and cancer (40, 41). Most clinical versions of monoclonal antibodies are now designed to have afucosylated Fc regions, so they are more capable of mediating ADCC and ADCP activities. As work continues toward developing a pan-CoV vaccine or therapy, it will be important to understand how to elicit optimal FcR functionality.

## MATERIALS AND METHODS

### Participants and Data Collection

Demographic and clinical information was extracted from the BMC EMR (Table S1). A representative subset of these individuals with available PBMCs were used in the T cell response analyses (Table S2). Pre-pandemic samples were collected from the HIV and aging cohort as previously described (42). The pre-pandemic samples were collected from June 2017 to March 2020, before the first diagnosed SARS-CoV-2 infected individual at BMC. All post-pandemic samples were collected after November 2020 and prior to December 2021 before the widespread Omicron SARS-CoV-2 surge (43). The post-pandemic blood samples were obtained from BMC patients during a non-COVID-19 related medical visit. Prior to sample collection, all individuals’ status regarding COVID-19 vaccination and prior SARS-CoV-2 infection was confirmed during the consent process. Previous SARS-CoV-2 infections were based on available prior SARS-CoV-2 reverse transcription (RT) -PCR test results. Vaccinated individuals had received at least two doses of the Pfizer BioNTech BNT162b2 or Moderna mRNA-1273 COVID-19 vaccine or one dose of the Janssen / Johnson & Johnson Ad26.COV2.S COVID-19 vaccine. No individual had received a COVID-19 vaccine booster because this practice was instituted after the end of our sample collection period. Included individuals were greater than 18 years of age. Documented eCoV infections were based on prior documented positive test results for HCoV-229E, HCoV-HKU1, HCoV-NL63, or HCoV-OC43 in the CRP-PCR test. CRP-PCR tests were used to evaluate patients that present with acute respiratory symptoms, but we did not confirm the medical reasons for the testing in all cases. The CRP-PCR was done at the discretion of the treating physician. This study was approved by the BMC Institutional Review Board.

### SARS-CoV-2-and ECoV-Specific IgG Antibody Quantification

SARS-CoV-2, eCoV spike, and nucleocapsid protein-specific IgG titers were detected by ELISA as previously described (25). SARS-CoV-2 RBD protein (SinoBiological, 40592-V08H), SARS-CoV-2 S2 protein (SinoBiological, 40590-V08H1), SARS-CoV-2 nucleocapsid protein (SinoBiological, 40588-V08B), HCoV-OC43 spike protein (SinoBiological, 40607-V08B), HCoV-OC43 nucleocapsid protein (SinoBiological, 40643-V07E), HCoV-229E spike protein (SinoBiological, 40605-V08B), HCoV-229E nucleocapsid protein (SinoBiological, 40640-V07E), HCoV-HKU1 nucleocapsid protein (SinoBiological, 40642-V07E), or HCoV-NL63 nucleocapsid protein (SinoBiological, 40641-V07E) were used in these ELISAs. After overnight incubation with an antigen, wells were blocked with casein blocking buffer (Thermo Fisher Scientific, 37528). Three plasma dilutions were tested (1:5, 1:100, 1:2000), and IgG was detected using anti-human horseradish peroxidase (HRP)-conjugated secondary antibodies for IgG detection (diluted 1:50000, Sigma-Aldrich, A0170) with 3,3’,5,5’ Tetramethylbenzidine (TMB)-ELISA substrate solution (Thermo Fisher Scientific, 34029). The reaction was stopped using 2M sulfuric acid and absorbance was measured on a SpectraMax190 Microplate Reader (Molecular Devices) at 450 nm. The optical density (OD) from the no antigen negative control wells was subtracted from all readings. A positive control standard (CR3022 IgG, Abcam, 273073) was serially diluted and measured against SARS-CoV-2 RBD to create a standard curve on each plate. The CR3022 standard curve was used to calculate titers (relative units) for each sample by interpolating a four-parameter logistic (4PL) curve.

### Pseudovirus Neutralization Assay

The vesicular stomatitis virus (VSV)-ΔG based pseudoviruses expressing SARS-CoV-2-Wuhan spikes were produced by transfecting SARS-CoV-2-spike protein (BEI Resources, NR52310) expression plasmids, infecting with (VSV)-G pseudotyped virus (G*ΔG-VSV), and collecting supernatants previously described (4). Pseudovirus neutralization assays were conducted as previously described (44). Briefly, heat inactivated plasma was serially diluted using a 2-fold serially dilution series starting at a 1:40 dilution and incubated with 1.25×10^4^ Vero E6 cells. Luciferase expression was measured using the Promega Bright-Glo Luciferase Assay System (Thermo Scientific) on a SpectraMax190 Microplate Reader (Molecular Devices). Percent neutralization was calculated in comparison to luciferase expression in infected wells without patient plasma. Area under the curve (AUC) values were calculated from the curve generated from the neutralizations across the serially diluted plasma (45). All neutralizations were tested in triplicate a minimum of two independent times. Neutralization against VSV-G pseudotyped virus (G*ΔG-VSV) was used as a control to assess activity against VSV-G protein in the absence of a CoV spike protein.

### SARS-CoV-2- and ECoV-Specific Antibody FcR Binding Quantification

Levels of SARS-CoV-2 and eCoV spike and nucleocapsid protein-specific antibodies with the ability to bind the Fc receptor, FcγRIIIa, were detected by the fucose-sensitive ELISA-based method of antigen-specific IgG Fc fucosylation (FEASI) (17). The ELISA protocol for antigen specific IgG quantification as described above (25) was adapted to include measurements of antibody binding to FcγRIIIa. The following additions were made to the previously described ELISA protocol. Plasma was diluted 1:5 to 1:50. Either 1ug/mL of His-tagged (Invitrogen, RP-87975) or biotinylated (gift from Dr. Gestur Vidarsson, PhD) FcγRIIIa was used as FcR antigens. Antibody bound FcR was detected by either anti-His HRP-conjugated antibody (1ug/mL, Invitrogen, MA1-21315-HRP) or HRP-conjugated streptavidin (0.125ug/mL, Thermo Scientific, N100). Similar methods of developing, reading, and calculating values for the ELISA were used as described earlier.

### SARS-CoV-2-Specific T cell Responses

The activation-induced marker (AIM) assay to measure antigen specific T cell responses was performed as described previously (46, 47). SARS-CoV-2 spike protein (BEI Resources, NR-52402), SARS-CoV-2 nucleocapsid (BEI Resources, NR-52404), SARS-CoV-2 nsp12 protein (JPT, PM-WCPV-NSP12-2), SARS-CoV-2 nsp13 protein (JPT, PM-WCPV-NSP13-2) pools were added at a final concentration of 1 µg/mL (containing less than 0.1% DMSO) were added for the stimulation. Media with 0.1% DMSO was used as a negative control.

After stimulation, the cells were fixed using cold BD Cytofix Fixation Buffer (1:10 dilution, BD Biosciences, 554655) and blocked using human Fc receptor block (1:100 dilution, BioLegend, 422302). Live cells were then stained for T cell lineage and activation markers: live/dead cell marker (1:200 dilution, Thermo Fisher, L23105), Alexa Fluor 647 anti-human CD3 (1:100 dilution, BioLegend, clone HIT3a, 300321), Alexa Fluor 488 anti-human CD4 (1:200 dilution, BioLegend, clone SK3, 344618), allophycocyanin (APC)/Fire 750 anti-human CD8a (1:50 dilution, BioLegend, clone HIT8a, 300931), phycoerythrin (PE)/Cyanine7 anti-human CD69 (1:50 dilution, BioLegend, clone FN50, 310911), Brilliant Violet 421 anti-human CD134 (OX40) (1:25 dilution, BioLegend, clone Ber-ACT35 (ACT-35), 350013), PE anti-human CD137 (4-1BB) (1:100 dilution, BioLegend, clone 4B4-1, 309803). Stained cells were analyzed on either a BD LSR II Flow Cytometer (BD Biosciences) or Cytek Aurora 5L (Cytek Biosciences).

The resulting flow cytometry data was analyzed using FlowJo software using similar methods as previously described (4). Live (Live/Dead marker ^-^) T cells (CD3^+^) were then gated on either CD4^+^ or CD8^+^. Activated CD4^+^ were defined by the double-positive CD134^+^ CD137^+^ population, while activated CD8^+^ were defined by the double-positive CD69^+^ CD137^+^ population. The percent of activated T cells for a given antigen stimulation condition was then background subtracted against the negative control (DMSO only) condition.

### Statistical Analysis

Individuals with a previous documented SARS-CoV-2 infection or previous COVID-19 vaccination were grouped together as those previous SARS-CoV-2 spike exposure for all spike-directed immune response assessments. Individuals with a previous COVID-19 vaccination were grouped based on their classified eCoV infection status for the non-spike directed immune response examinations. Composite alpha or beta eCoV nucleocapsid antibody scores were created by Log2 transforming the average of HCoV-229E and HCoV-NL63 or HCoV-HKU1 and HCoV-OC43 IgG titers, respectively. Individuals with either a documented eCoV infection in the past 550 days or those within the top 25% with the highest alpha or beta eCoV nucleocapsid antibody scores were classified as presumed or documented eCoV infection. The remaining SARS-CoV-2 infection naïve individuals were considered without a recent eCoV infection.

Comparison of responses in individuals with a previous SARS-CoV-2 infection or SARS-CoV-2 spike exposure to the SARS-CoV-2 infection naïve groups was conducted using Kruskal-Wallis tests and corrected with Dunn’s multiple comparisons test. Mann-Whitney U tests were used for direct comparisons between individuals with or without a presumed recent eCoV infection. Correlations were assessed using Pearson and Spearman tests as appropriate. Categorical differences were examined using Fischer’s exact test or Chi-square test when comparing more than 2 groups. In the multivariable logistic regression model, the presence or absence of a SARS-CoV-2 antibody binding FcγRIIIa response was the dependent variable. Alpha and beta eCoV nucleocapsid IgG levels, prior SARS-CoV-2 spike experience (categorical variable), and available demographic factors and co-morbidities were independent predictors in the multivariate analysis. All covariates in a univariate model with p values less or equal to 0.15 level were initially included in the multivariable model. Covariates with p values greater than 0.10 in the multivariable model were then removed. Statistical analyses were performed using GraphPad Prism 9.0.2. A two-sided p-value less than 0.05 was considered statistically significant.

## ACKNOWLEDGMENTS

We thank the study participants for providing information and donating specimens. We thank the Boston University Flow Cytometry Core. The content is solely the responsibility of the authors and does not necessarily represent the official views of the National Institutes of Health. This study was supported by National Institutes of Health grants including K24-AI145661 and P30-AI042853. DJB was funded through a T32-5T32AI00730928. Massachusetts Consortium on Pathogen Readiness funded sample collection.

## REFERENCES

1. Hamre D, Beem M. 1972. Virologic studies of acute respiratory disease in young adults. V. Coronavirus 229E infections during six years of surveillance. Am J Epidemiol 96:94–106.

2. Edridge AWD, Kaczorowska J, Hoste ACR, Bakker M, Klein M, Loens K, Jebbink MF, Matser A, Kinsella CM, Rueda P, Ieven M, Goossens H, Prins M, Sastre P, Deijs M, van der Hoek L. 2020. Seasonal coronavirus protective immunity is short-lasting. Nat Med 26:1691–1693.

3. Dangi T, Palacio N, Sanchez S, Park M, Class J, Visvabharathy L, Ciucci T, Koralnik IJ, Richner JM, Penaloza-MacMaster P. 2021. Cross-protective immunity following coronavirus vaccination and coronavirus infection. J Clin Invest 131:e151969.

4. Bean DJ, Monroe J, Liang YM, Borberg E, Senussi Y, Swank Z, Chalise S, Walt D, Weinberg J, Sagar M. 2024. Heterotypic immunity from prior SARS-CoV-2 infection but not COVID-19 vaccination associates with lower endemic coronavirus incidence. Sci Transl Med 16:eado7588.

5. Sagar M, Reifler K, Rossi M, Miller NS, Sinha P, White LF, Mizgerd JP. 2021. Recent endemic coronavirus infection is associated with less-severe COVID-19. J Clin Invest 131:e143380.

6. Swadling L, Diniz MO, Schmidt NM, Amin OE, Chandran A, Shaw E, Pade C, Gibbons JM, Le Bert N, Tan AT, Jeffery-Smith A, Tan CCS, Tham CYL, Kucykowicz S, Aidoo-Micah G, Rosenheim J, Davies J, Johnson M, Jensen MP, Joy G, McCoy LE, Valdes AM, Chain BM, Goldblatt D, Altmann DM, Boyton RJ, Manisty C, Treibel TA, Moon JC, van Dorp L, Balloux F, McKnight Á, Noursadeghi M, Bertoletti A, Maini MK. 2022. Pre-existing polymerase-specific T cells expand in abortive seronegative SARS-CoV-2. Nature 601:110–117.

7. Loyal L, Braun J, Henze L, Kruse B, Dingeldey M, Reimer U, Kern F, Schwarz T, Mangold M, Unger C, Dörfler F, Kadler S, Rosowski J, Gürcan K, Uyar-Aydin Z, Frentsch M, Kurth F, Schnatbaum K, Eckey M, Hippenstiel S, Hocke A, Müller MA, Sawitzki B, Miltenyi S, Paul F, Mall MA, Wenschuh H, Voigt S, Drosten C, Lauster R, Lachman N, Sander L-E, Corman VM, Röhmel J, Meyer-Arndt L, Thiel A, Giesecke-Thiel C. 2021. Cross-reactive CD4+ T cells enhance SARS-CoV-2 immune responses upon infection and vaccination. Science 374:eabh1823.

8. Ortega N, Ribes M, Vidal M, Rubio R, Aguilar R, Williams S, Barrios D, Alonso S, Hernández-Luis P, Mitchell RA, Jairoce C, Cruz A, Jimenez A, Santano R, Méndez S, Lamoglia M, Rosell N, Llupià A, Puyol L, Chi J, Melero NR, Parras D, Serra P, Pradenas E, Trinité B, Blanco J, Mayor A, Barroso S, Varela P, Vilella A, Trilla A, Santamaria P, Carolis C, Tortajada M, Izquierdo L, Angulo A, Engel P, García-Basteiro AL, Moncunill G, Dobaño C. 2021. Seven-month kinetics of SARS-CoV-2 antibodies and role of pre-existing antibodies to human coronaviruses. Nat Commun 12:4740.

9. Walsh EE, Shin JH, Falsey AR. 2013. Clinical Impact of Human Coronaviruses 229E and OC43 Infection in Diverse Adult Populations. The Journal of Infectious Diseases 208:1634–1642.

10. Gaunt ER, Hardie A, Claas ECJ, Simmonds P, Templeton KE. 2010. Epidemiology and clinical presentations of the four human coronaviruses 229E, HKU1, NL63, and OC43 detected over 3 years using a novel multiplex real-time PCR method. J Clin Microbiol 48:2940–2947.

11. Leber AL, Everhart K, Daly JA, Hopper A, Harrington A, Schreckenberger P, McKinley K, Jones M, Holmberg K, Kensinger B. 2018. Multicenter Evaluation of BioFire FilmArray Respiratory Panel 2 for Detection of Viruses and Bacteria in Nasopharyngeal Swab Samples. Journal of Clinical Microbiology 56:10.1128/jcm.01945-17.

12. Burbelo PD, Riedo FX, Morishima C, Rawlings S, Smith D, Das S, Strich JR, Chertow DS, Davey RT Jr, Cohen JI. 2020. Sensitivity in Detection of Antibodies to Nucleocapsid and Spike Proteins of Severe Acute Respiratory Syndrome Coronavirus 2 in Patients With Coronavirus Disease 2019. The Journal of Infectious Diseases 222:206–213.

13. Blanchard EG, Miao C, Haupt TE, Anderson LJ, Haynes LM. 2011. Development of a recombinant truncated nucleocapsid protein based immunoassay for detection of antibodies against human coronavirus OC43. J Virol Methods 177:100–106.

14. Garcia-Beltran WF, Lam EC, Astudillo MG, Yang D, Miller TE, Feldman J, Hauser BM, Caradonna TM, Clayton KL, Nitido AD, Murali MR, Alter G, Charles RC, Dighe A, Branda JA, Lennerz JK, Lingwood D, Schmidt AG, Iafrate AJ, Balazs AB. 2021. COVID-19-neutralizing antibodies predict disease severity and survival. Cell 184:476–488.e11.

15. Khoury DS, Cromer D, Reynaldi A, Schlub TE, Wheatley AK, Juno JA, Subbarao K, Kent SJ, Triccas JA, Davenport MP. 2021. Neutralizing antibody levels are highly predictive of immune protection from symptomatic SARS-CoV-2 infection. Nat Med 27:1205–1211.

16. Yu Y, Wang M, Zhang X, Li S, Lu Q, Zeng H, Hou H, Li H, Zhang M, Jiang F, Wu J, Ding R, Zhou Z, Liu M, Si W, Zhu T, Li H, Ma J, Gu Y, She G, Li X, Zhang Y, Peng K, Huang W, Liu W, Wang Y. 2021. Antibody-dependent cellular cytotoxicity response to SARS-CoV-2 in COVID-19 patients. Signal Transduct Target Ther 6:346.

17. Šuštić T, Van Coillie J, Larsen MD, Derksen NIL, Szittner Z, Nouta J, Wang W, Damelang T, Rebergen I, Linty F, Visser R, Mok JY, Geerdes DM, van Esch WJE, de Taeye SW, van Gils MJ, van de Watering L, van der Schoot CE, Wuhrer M, Rispens T, Vidarsson G. 2022. Immunoassay for quantification of antigen-specific IgG fucosylation. EBioMedicine 81:104109.

18. Peiris J, Lai S, Poon L, Guan Y, Yam L, Lim W, Nicholls J, Yee W, Yan W, Cheung M, Cheng V, Chan K, Tsang D, Yung R, Ng T, Yuen K. 2003. Coronavirus as a possible cause of severe acute respiratory syndrome. Lancet 361:1319–1325.

19. de Groot RJ, Baker SC, Baric RS, Brown CS, Drosten C, Enjuanes L, Fouchier RAM, Galiano M, Gorbalenya AE, Memish ZA, Perlman S, Poon LLM, Snijder EJ, Stephens GM, Woo PCY, Zaki AM, Zambon M, Ziebuhr J. 2013. Middle East Respiratory Syndrome Coronavirus (MERS-CoV): Announcement of the Coronavirus Study Group. J Virol 87:7790–7792.

20. Zhou P, Yang X-L, Wang X-G, Hu B, Zhang L, Zhang W, Si H-R, Zhu Y, Li B, Huang C-L, Chen H-D, Chen J, Luo Y, Guo H, Jiang R-D, Liu M-Q, Chen Y, Shen X-R, Wang X, Zheng X-S, Zhao K, Chen Q-J, Deng F, Liu L-L, Yan B, Zhan F-X, Wang Y-Y, Xiao G-F, Shi Z-L. 2020. A pneumonia outbreak associated with a new coronavirus of probable bat origin. Nature 579:270–273.

21. Abela IA, Schwarzmüller M, Ulyte A, Radtke T, Haile SR, Ammann P, Raineri A, Rueegg S, Epp S, Berger C, Böni J, Manrique A, Audigé A, Huber M, Schreiber PW, Scheier T, Fehr J, Weber J, Rusert P, Günthard HF, Kouyos RD, Puhan MA, Kriemler S, Trkola A, Pasin C. 2024. Cross-protective HCoV immunity reduces symptom development during SARS-CoV-2 infection. mBio 15:e02722–23.

22. Aguilar-Bretones M, Westerhuis BM, Raadsen MP, Bruin E de, Chandler FD, Okba NMA, Haagmans BL, Langerak T, Endeman H, Akker JPC van den, Gommers DAMPJ, Gorp ECM van, GeurtsvanKessel CH, Vries RD de, Fouchier RAM, Rockx BHG, Koopmans MPG, Nierop GP van. 2021. Seasonal coronavirus–specific B cells with limited SARS-CoV-2 cross-reactivity dominate the IgG response in severe COVID-19. J Clin Invest 131.

23. Anderson EM, Goodwin EC, Verma A, Arevalo CP, Bolton MJ, Weirick ME, Gouma S, McAllister CM, Christensen SR, Weaver J, Hicks P, Manzoni TB, Oniyide O, Ramage H, Mathew D, Baxter AE, Oldridge DA, Greenplate AR, Wu JE, Alanio C, D’Andrea K, Kuthuru O, Dougherty J, Pattekar A, Kim J, Han N, Apostolidis SA, Huang AC, Vella LA, Kuri-Cervantes L, Pampena MB, Betts MR, Wherry EJ, Meyer NJ, Cherry S, Bates P, Rader DJ, Hensley SE. 2021. Seasonal human coronavirus antibodies are boosted upon SARS-CoV-2 infection but not associated with protection. Cell 184:1858–1864.e10.

24. Mateus J, Grifoni A, Tarke A, Sidney J, Ramirez SI, Dan JM, Burger ZC, Rawlings SA, Smith DM, Phillips E, Mallal S, Lammers M, Rubiro P, Quiambao L, Sutherland A, Yu ED, da Silva Antunes R, Greenbaum J, Frazier A, Markmann AJ, Premkumar L, de Silva A, Peters B, Crotty S, Sette A, Weiskopf D. 2020. Selective and cross-reactive SARS-CoV-2 T cell epitopes in unexposed humans. Science 370:89–94.

25. Yuen RR, Steiner D, Pihl RMF, Chavez E, Olson A, Smith EL, Baird LA, Korkmaz F, Urick P, Sagar M, Berrigan JL, Gummuluru S, Corley RB, Quillen K, Belkina AC, Mostoslavsky G, Rifkin IR, Kataria Y, Cappione AJ, Gao W, Lin NH, Bhadelia N, Snyder-Cappione JE. 2021. Novel ELISA Protocol Links Pre-Existing SARS-CoV-2 Reactive Antibodies With Endemic Coronavirus Immunity and Age and Reveals Improved Serologic Identification of Acute COVID-19 via Multi-Parameter Detection. Front Immunol 12:614676.

26. Dijkman R, Jebbink MF, El Idrissi NB, Pyrc K, Müller MA, Kuijpers TW, Zaaijer HL, van der Hoek L. 2008. Human coronavirus NL63 and 229E seroconversion in children. J Clin Microbiol 46:2368–2373.

27. Grobben M, van der Straten K, Brouwer PJ, Brinkkemper M, Maisonnasse P, Dereuddre-Bosquet N, Appelman B, Lavell AA, van Vught LA, Burger JA, Poniman M, Oomen M, Eggink D, Bijl TP, van Willigen HD, Wynberg E, Verkaik BJ, Figaroa OJ, de Vries PJ, Boertien TM, Amsterdam UMC COVID-19 S3/HCW study group, Bomers MK, Sikkens JJ, Le Grand R, de Jong MD, Prins M, Chung AW, de Bree GJ, Sanders RW, van Gils MJ. 2021. Cross-reactive antibodies after SARS-CoV-2 infection and vaccination. Elife 10:e70330.

28. Geanes ES, LeMaster C, Fraley ER, Khanal S, McLennan R, Grundberg E, Selvarangan R, Bradley T. 2022. Cross-reactive antibodies elicited to conserved epitopes on SARS-CoV-2 spike protein after infection and vaccination. Sci Rep 12:6496.

29. Adeniji OS, Giron LB, Purwar M, Zilberstein NF, Kulkarni AJ, Shaikh MW, Balk RA, Moy JN, Forsyth CB, Liu Q, Dweep H, Kossenkov A, Weiner DB, Keshavarzian A, Landay A, Abdel-Mohsen M. 2021. COVID-19 Severity Is Associated with Differential Antibody Fc-Mediated Innate Immune Functions. mBio 12:e00281–21.

30. Maucourant C, Filipovic I, Ponzetta A, Aleman S, Cornillet M, Hertwig L, Strunz B, Lentini A, Reinius B, Brownlie D, Cuapio A, Ask EH, Hull RM, Haroun-Izquierdo A, Schaffer M, Klingström J, Folkesson E, Buggert M, Sandberg JK, Eriksson LI, Rooyackers O, Ljunggren H-G, Malmberg K-J, Michaëlsson J, Marquardt N, Hammer Q, Strålin K, Björkström NK, Karolinska COVID-19 Study Group. 2020. Natural killer cell immunotypes related to COVID-19 disease severity. Sci Immunol 5:eabd6832.

31. Kiyoshi M, Caaveiro JMM, Kawai T, Tashiro S, Ide T, Asaoka Y, Hatayama K, Tsumoto K. 2015. Structural basis for binding of human IgG1 to its high-affinity human receptor FcγRI. Nat Commun 6:6866.

32. Chakraborty S, Gonzalez JC, Sievers BL, Mallajosyula V, Chakraborty S, Dubey M, Ashraf U, Cheng BY-L, Kathale N, Tran KQT, Scallan C, Sinnott A, Cassidy A, Chen ST, Gelbart T, Gao F, Golan Y, Ji X, Kim-Schulze S, Prahl M, Gaw SL, Gnjatic S, Marron TU, Merad M, Arunachalam PS, Boyd SD, Davis MM, Holubar M, Khosla C, Maecker HT, Maldonado Y, Mellins ED, Nadeau KC, Pulendran B, Singh U, Subramanian A, Utz PJ, Sherwood R, Zhang S, Jagannathan P, Tan GS, Wang TT. 2022. Early non-neutralizing, afucosylated antibody responses are associated with COVID-19 severity. Science Translational Medicine 14:eabm7853.

33. Larsen MD, de Graaf EL, Sonneveld ME, Plomp HR, Nouta J, Hoepel W, Chen H-J, Linty F, Visser R, Brinkhaus M, Šuštić T, de Taeye SW, Bentlage AEH, Toivonen S, Koeleman CAM, Sainio S, Kootstra NA, Brouwer PJM, Geyer CE, Derksen NIL, Wolbink G, de Winther M, Sanders RW, van Gils MJ, de Bruin S, Vlaar APJ, Rispens T, den Dunnen J, Zaaijer HL, Wuhrer M, Ellen van der Schoot C, Vidarsson G. 2021. Afucosylated IgG characterizes enveloped viral responses and correlates with COVID-19 severity. Science 371:eabc8378.

34. Junqueira C, Crespo Â, Ranjbar S, de Lacerda LB, Lewandrowski M, Ingber J, Parry B, Ravid S, Clark S, Schrimpf MR, Ho F, Beakes C, Margolin J, Russell N, Kays K, Boucau J, Das Adhikari U, Vora SM, Leger V, Gehrke L, Henderson LA, Janssen E, Kwon D, Sander C, Abraham J, Goldberg MB, Wu H, Mehta G, Bell S, Goldfeld AE, Filbin MR, Lieberman J. 2022. FcγR-mediated SARS-CoV-2 infection of monocytes activates inflammation. Nature 606:576–584.

35. Bergamaschi L, Mescia F, Turner L, Hanson AL, Kotagiri P, Dunmore BJ, Ruffieux H, De Sa A, Huhn O, Morgan MD, Gerber PP, Wills MR, Baker S, Calero-Nieto FJ, Doffinger R, Dougan G, Elmer A, Goodfellow IG, Gupta RK, Hosmillo M, Hunter K, Kingston N, Lehner PJ, Matheson NJ, Nicholson JK, Petrunkina AM, Richardson S, Saunders C, Thaventhiran JED, Toonen EJM, Weekes MP, Cambridge Institute of Therapeutic Immunology and Infectious Disease-National Institute of Health Research (CITIID-NIHR) COVID BioResource Collaboration, Göttgens B, Toshner M, Hess C, Bradley JR, Lyons PA, Smith KGC. 2021. Longitudinal analysis reveals that delayed bystander CD8+ T cell activation and early immune pathology distinguish severe COVID-19 from mild disease. Immunity 54:1257–1275.e8.

36. Yu M, Charles A, Cagigi A, Christ W, Österberg B, Falck-Jones S, Azizmohammadi L, Åhlberg E, Falck-Jones R, Svensson J, Nie M, Warnqvist A, Hellgren F, Lenart K, Arcoverde Cerveira R, Ols S, Lindgren G, Lin A, Maecker H, Bell M, Johansson N, Albert J, Sundling C, Czarnewski P, Klingström J, Färnert A, Loré K, Smed-Sörensen A. 2023. Delayed generation of functional virus-specific circulating T follicular helper cells correlates with severe COVID-19. Nat Commun 14:2164.

37. Low JS, Vaqueirinho D, Mele F, Foglierini M, Jerak J, Perotti M, Jarrossay D, Jovic S, Perez L, Cacciatore R, Terrot T, Pellanda AF, Biggiogero M, Garzoni C, Ferrari P, Ceschi A, Lanzavecchia A, Sallusto F, Cassotta A. 2021. Clonal analysis of immunodominance and cross-reactivity of the CD4 T cell response to SARS-CoV-2. Science 372:1336–1341.

38. Bacher P, Rosati E, Esser D, Martini GR, Saggau C, Schiminsky E, Dargvainiene J, Schröder I, Wieters I, Khodamoradi Y, Eberhardt F, Vehreschild MJGT, Neb H, Sonntagbauer M, Conrad C, Tran F, Rosenstiel P, Markewitz R, Wandinger K-P, Augustin M, Rybniker J, Kochanek M, Leypoldt F, Cornely OA, Koehler P, Franke A, Scheffold A. 2020. Low-Avidity CD4+ T Cell Responses to SARS-CoV-2 in Unexposed Individuals and Humans with Severe COVID-19. Immunity 53:1258–1271.e5.

39. Tarke A, Sidney J, Kidd CK, Dan JM, Ramirez SI, Yu ED, Mateus J, da Silva Antunes R, Moore E, Rubiro P, Methot N, Phillips E, Mallal S, Frazier A, Rawlings SA, Greenbaum JA, Peters B, Smith DM, Crotty S, Weiskopf D, Grifoni A, Sette A. 2021. Comprehensive analysis of T cell immunodominance and immunoprevalence of SARS-CoV-2 epitopes in COVID-19 cases. Cell Rep Med 2:100204.

40. Thomas AS, Moreau Y, Jiang W, Isaac JE, Ewing A, White LF, Kourtis AP, Sagar M. 2021. Pre-existing infant antibody-dependent cellular cytotoxicity associates with reduced HIV-1 acquisition and lower morbidity. Cell Rep Med 2:100412.

41. Junttila TT, Parsons K, Olsson C, Lu Y, Xin Y, Theriault J, Crocker L, Pabonan O, Baginski T, Meng G, Totpal K, Kelley RF, Sliwkowski MX. 2010. Superior in vivo efficacy of afucosylated trastuzumab in the treatment of HER2-amplified breast cancer. Cancer Res 70:4481–4489.

42. Belkina AC, Starchenko A, Drake KA, Proctor EA, Pihl RMF, Olson A, Lauffenburger DA, Lin N, Snyder-Cappione JE. 2018. Multivariate Computational Analysis of Gamma Delta T Cell Inhibitory Receptor Signatures Reveals the Divergence of Healthy and ART-Suppressed HIV+ Aging. Front Immunol 9:2783.

43. Clarke KEN, Jones JM, Deng Y, Nycz E, Lee A, Iachan R, Gundlapalli AV, Hall AJ, MacNeil A. 2022. Seroprevalence of Infection-Induced SARS-CoV-2 Antibodies - United States, September 2021-February 2022. MMWR Morb Mortal Wkly Rep 71:606–608.

44. Nie J, Li Q, Wu J, Zhao C, Hao H, Liu H, Zhang L, Nie L, Qin H, Wang M, Lu Q, Li X, Sun Q, Liu J, Fan C, Huang W, Xu M, Wang Y. 2020. Quantification of SARS-CoV-2 neutralizing antibody by a pseudotyped virus-based assay. Nat Protoc 15:3699–3715.

45. Yu X, Gilbert PB, Hioe CE, Zolla-Pazner S, Self SG. 2012. Statistical approaches to analyzing HIV-1 neutralizing antibody assay data. Stat Biopharm Res 4:1–13.

46. Dan JM, Lindestam Arlehamn CS, Weiskopf D, da Silva Antunes R, Havenar-Daughton C, Reiss SM, Brigger M, Bothwell M, Sette A, Crotty S. 2016. A Cytokine-Independent Approach To Identify Antigen-Specific Human Germinal Center T Follicular Helper Cells and Rare Antigen-Specific CD4+ T Cells in Blood. The Journal of Immunology 197:983–993.

47. Reiss S, Baxter AE, Cirelli KM, Dan JM, Morou A, Daigneault A, Brassard N, Silvestri G, Routy J-P, Havenar-Daughton C, Crotty S, Kaufmann DE. 2017. Comparative analysis of activation induced marker (AIM) assays for sensitive identification of antigen-specific CD4 T cells. PLoS One 12:e0186998.

