## Supplemental Files for "Recent Endemic Coronavirus Infection Associates With Higher SARS-CoV-2 Cross-Reactive Fc Receptor Binding Antibodies"

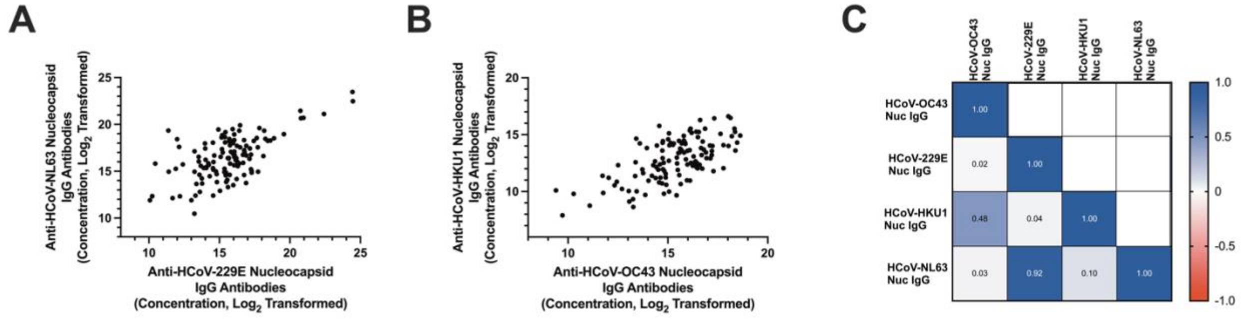


**Figure S1. Correlations between IgG antibody responses to alpha and beta eCoV nucleocapsid proteins.** Anti-nucleocapsid antibody responses were measured in all individuals in the study. (A) Correlation of IgG antibody responses against HCoV-229E and HCoV-NL63 nucleocapsids. (B) Correlation of IgG antibody responses against HCoV-HKU1 and HCoV-OC43 nucleocapsids. (C) The matrix shows Pearson correlation r values between IgG antibody responses against HCoV-OC43, HCoV-229E, HCoV-HKU1, and HCoV-NL63 nucleocapsid proteins. Blue represents stronger positive correlations, while red signifies stronger negative correlations.

**Table S1. Demographics of the individuals with collected blood specimens for analyses of SARS-CoV-2 and eCoV mediated antibody responses**

| | Prior SARS-CoV-2 infection / No COVID-19 vaccine (n = 20) | Prior COVID-19 vaccine / No SARS-CoV-2 infection (n = 29) | No prior SARS-CoV-2 infection or COVID-19 vaccine (n = 72) | p-value [a] |
|---|---|---|---|---|
| **Age (years), median (IQR)** | 53 (48 – 59) | 62 (54 – 70) | 56 (46 – 64) | 0.0215 [b] |
| **Male** | 8 (40) | 19 (66) | 36 (50) | 0.1834 |
| **Race/Ethnicity [c]** | | | | 0.3087 |
| **Black** | 14 (70) | 13 (45) | 35 (49) | |
| **White** | 3 (15) | 13 (45) | 26 (36) | |
| **Hispanic/Latino** | 3 (15) | 3 (10) | 8 (11) | |
| **Other/missing** | 0 (0) | 0 (0) | 3 (4) | |
| **Diabetes mellitus** | 5 (25) | 10 (34) | 21 (29) | 0.7640 |
| **Heart disease [d]** | 3 (15) | 12 (41) | 15 (21) | 0.0520 |
| **Lung disease [e]** | 5 (25) | 8 (28) | 26 (36) | 0.5322 |
| **CKD [f]** | 1 (5) | 3 (10) | 6 (8) | 0.7997 |
| **HIV [g]** | 7 (35) | 5 (17) | 26 (36) | 0.1686 |
| **Cancer** | 1 (5) | 2 (7) | 1 (1) | 0.3367 |
| **Number of co-morbidities [h]** | | | | 0.6309 |
| **0** | 3 (15) | 5 (17) | 12 (17) | |
| **1** | 12 (60) | 12 (41) | 33 (46) | |
| **≥2** | 5 (25) | 12 (41) | 17 (38) | |
| **Pre-Pandemic [i]** | - | - | 14 (26) | - |

Data shows number and percent unless otherwise indicated, [a] Chi-square test unless otherwise indicated, [b] Kruskal-Wallis test and Dunn's multiple comparison test, [c] As specified in the EMR; an individual may be in more than 1 category, [d] Heart disease includes coronary artery disease and congestive heart failure, [e] Lung disease includes chronic obstructive pulmonary disease and asthma, [f] Chronic kidney disease, [g] Human immunodeficiency virus, [h] Number of comorbidities accounts for diabetes mellitus, heart disease, lung disease, chronic kidney disease, HIV, and cancer, [i] Samples collected before March 2020

**Table S2. Demographics of the individuals with available PBMCs for analyses of eCoV mediated cellular responses against SARS-CoV-2 antigens**

| | Prior SARS-CoV-2 infection / No COVID-19 vaccine (n = 20) | Presumed or documented recent eCoV infection (n = 32) | No presumed recent eCoV infection (n = 42) | p-value [a] |
|---|---|---|---|---|
| **Age (years), median (IQR)** | 53 (48 – 59) | 65 (57 – 71) | 57 (44 – 62) | 0.0057 [b] |
| **Male** | 8 (40) | 17 (53) | 22 (52) | 0.6004 |
| **Race/Ethnicity [c]** | | | | 0.3752 |
| **Black** | 14 (70) | 12 (38) | 21 (50) | |
| **White** | 3 (15) | 14 (44) | 15 (36) | |
| **Hispanic/Latino** | 3 (15) | 5 (16) | 5 (12) | |
| **Other/missing** | 0 (0) | 1 (3) | 1 (2) | |
| **Diabetes mellitus** | 5 (25) | 12 (38) | 15 (36) | 0.6216 |
| **Heart disease [d]** | 3 (15) | 13 (41) | 9 (21) | 0.0751 |
| **Lung disease [e]** | 5 (25) | 13 (41) | 8 (19) | 0.1156 |
| **CKD [f]** | 1 (5) | 2 (6) | 5 (12) | 0.5633 |
| **HIV [g]** | 7 (35) | 8 (25) | 14 (33) | 0.6716 |
| **Cancer** | 1 (5) | 1 (3) | 1 (2) | 0.8601 |
| **Number of co-morbidities [h]** | | | | 0.3873 |
| **0** | 3 (15) | 5 (16) | 8 (19) | |
| **1** | 12 (60) | 11 (34) | 19 (45) | |
| **≥2** | 5 (25) | 16 (50) | 15 (36) | |
| **Pre-Pandemic [i]** | - | 4 (13) | 14 (33) | 0.0553 [j] |
| **COVID-19 Vaccine** | - | 13 (41) | 10 (24) | 0.1368 [j] |

Data shows number and percent unless otherwise indicated, [a] Chi-square test unless otherwise indicated, [b] Kruskal-Wallis test and Dunn's multiple comparison test, [c] As specified in the EMR; an individual may be in more than 1 category, [d] Heart disease includes coronary artery disease and congestive heart failure, [e] Lung disease includes chronic obstructive pulmonary disease and asthma, [f] Chronic kidney disease, [g] Human immunodeficiency virus, [h] Number of comorbidities accounts for diabetes mellitus, heart disease, lung disease, chronic kidney disease, HIV, and cancer, [i] Samples collected before March 2020, [j] Fisher's Exact Test

**Table S3. Statistical comparison of SARS-CoV-2 S2-specific antibody FcγRIIIa binding responses between individuals with or without a presumed recent eCoV infection**

| Percent of individuals with highest anti-alpha and beta eCoV nucleocapsids IgG levels included in the presumed recent eCoV group | Presumed or documented recent eCoV infection (median, IQR) | No presumed recent eCoV infection (median, IQR) | p-value [a] |
|---|---|---|---|
| 25% | 1.662 (0.587-3.165) | 0.100 (0.100-0.977) | 0.0234 |
| 30% | 1.436 (0.280-2.369) | 0.100 (0.100-0.977) | 0.0629 |
| 35% | 1.071 (0.145-2.298) | 0.100 (0.100-1.021) | 0.1037 |

[a] Mann-Whitney test between groups with or without a presumed recent eCoV infection
